# The chromatin remodeller CHD4 regulates transcription factor binding to both prevent activation of silent enhancers and maintain active regulatory elements

**DOI:** 10.1101/2025.08.29.672645

**Authors:** Andria Koulle, Oluwaseun Ogundele, Devina Shah, India-May Baker, Maya Lopez, Nicola Reynolds, Ramy Ragheb, Ernest Laue, Brian Hendrich

## Abstract

Chromatin organisation and transcriptional regulation are tightly coordinated processes that are essential for maintaining cellular identity and function. ATP-dependent chromatin remodelling proteins play critical roles in control of genome structure and in regulating transcription across eukaryotes. Their essential nature, however, has made it difficult to define exactly how these functions are mediated. The chromatin remodeller CHD4 has been shown to be capable of sliding nucleosomes in vitro, and to regulate chromatin accessibility and gene expression in vivo. Using an inducible depletion system, here we identify a second mechanism of action for CHD4 in actively restricting the residence time of transcription factors on chromatin. Together these activities result in distinct, context-dependent outcomes: at highly accessible regulatory elements, CHD4 limits transcription factor binding to maintain regulatory function, while at low-accessibility euchromatic regions, it prevents transcription factor engagement and sustains chromatin compaction, thereby silencing cryptic enhancers. Collectively, these mechanisms enable CHD4 to reduce transcriptional noise while preserving the responsiveness of active regulatory networks.

## Introduction

Cell state transitions are driven by the activity of transcription factors. Many transcription factors bind to specific DNA motifs within chromatin, but the frequency of these motifs within mammalian genomes far outnumbers sites at which protein binding is detectable. The ability of a transcription factor to recognise its cognate motif is influenced by how the DNA encoding that motif is packaged in chromatin. Accessible sites, i.e. those associated with a low density of intact nucleosomes, are more likely to be identified and bound by transcription factors than are inaccessible sites, i.e. those associated with higher nucleosome density. Controlling chromatin accessibility is therefore crucial for controlling transcription factor binding to cognate sites in regulatory regions. Transcription factor binding patterns will then determine which gene regulatory regions are used to drive gene expression and thereby define cell identity.

Vertebrate cells contain multiple proteins capable of using energy derived from ATP hydrolysis to remodel nucleosomes. These chromatin remodellers share a conserved sucrose non-fermentable 2 (SNF2) helicase-like ATPase domain but otherwise have various additional functional domains which impact how they organise chromatin, transcription and DNA repair (Hota and Bruneau, 2016; Narlikar et al., 2013). Chromatin remodellers play important roles in mammalian development, and heterozygous mutations in the genes encoding them underlie a variety of developmental disorders in humans (Gourisankar et al., 2024; Hota and Bruneau, 2016; Pierson et al., 2019). Similarly, somatic mutations in chromatin remodeller subunit genes are increasingly being implicated in cancer initiation and/or progression (Lai and Wade, 2011; Martincorena et al., 2017; Wilson and Roberts, 2011).

CHD4 is an abundant ATP-dependent chromatin remodelling protein which plays important roles in chromatin organisation and cell fate decisions in many different aspects of metazoan development. Depletion of CHD4 in mouse or Drosophila cells, or overexpression of a dominant negative form of the protein, leads to increased chromatin accessibility at regulatory elements and DNase hypersensitive sites (Dieuleveult et al., 2016; Morris et al., 2014; Moshkin et al., 2012), indicating that it predominantly functions to compact chromatin. Its impact on gene expression is less clear-cut, however. One study found that while CHD4 acted to repress genes with bivalent promoters, its activity at active promoters (marked with H3K4Me3) mainly facilitated transcription in mouse embryonic stem cells (Dieuleveult et al., 2016). Another study found CHD4 controlled the probability of gene expression, rather than levels, during the first cell fate transition in mammalian embryogenesis (O’Shaughnessy-Kirwan et al., 2015). Several studies in somatic lineages have shown that CHD4 prevents lineage inappropriate gene expression during cell fate decisions in both mice and Drosophila (Arends et al., 2019; Aughey et al., 2023; Gómez-del Arco et al., 2016; Sreenivasan et al., 2021; Wilczewski et al., 2018; Yoshida et al., 2019). Exactly how CHD4 has this varied impact on chromatin and transcription has not been defined.

CHD4 is the predominant chromatin remodelling subunit of the Nucleosome Remodelling and Deacetylation (NuRD) complex (Wade et al., 1998; Xue et al., 1998; Zhang et al., 1998). NuRD is a highly abundant chromatin remodeller present at active enhancers and promoters in many cell types. Its activity is important not only to regulate transcription but also the movement of enhancers in 3D space (Basu et al., 2023), and to maintain genome integrity (Polo et al., 2010; Smeenk et al., 2010). NuRD has been shown to exert two functions in mammalian cells: one is to control the nucleosome density at active enhancers, thereby regulating transcription factor binding, enhancer activity and transcriptional output (Bornelöv et al., 2018; Pundhir et al., 2023), while the other is to prevent low-level, inappropriate transcription across the genome (Burgold et al., 2019; Montibus et al., 2023; Ragheb et al., 2020; Saotome et al., 2024).

Although mouse ES cells can survive with a complete loss of the histone deacetylase subunit of NuRD (Burgold et al., 2019), loss of the chromatin remodelling component, CHD4, leads to cell death (Stevens et al., 2017). While we have used genetics to create NuRD-low or NuRD-null ES cells in the past to define NuRD function, in both cases, the remodelling subcomplex (CHD4, GATAD2A/B, CDK2AP1) remained on chromatin and the extent to which it could continue to remodel chromatin is not known (Bornelöv et al., 2018). As with any genetic change, it is also very difficult to know to what extent constitutive loss of protein activity has resulted in selection of cells using some compensatory mechanism to remain viable. CHD4 is also known to function outside of the NuRD complex (O’Shaughnessy-Kirwan et al., 2015; Ostapcuk et al., 2018; Williams et al., 2004). Assessing the direct function of NuRD’s remodelling activity is therefore difficult as knockout of the remodeller results in a non-viable cell, while knockdown or exogenous overexpression of a dominant negative version of the remodeller (Bornelöv et al., 2018; Feng and Zhang, 2001) will likely produce a heterogeneous mix of cells with varying levels of remodeller activity and displaying increasing degrees of cell cycle arrest and apoptosis over time.

The CHD4/NuRD function described at enhancers has been defined both in genetic mutants and over a time course of NuRD reintroduction to null cells (Bornelöv et al., 2018; Reynolds et al., 2012). In contrast, the noise reduction function has been described in genetically NuRD-deficient or NuRD-null cells or after NuRD component knockdown (Burgold et al., 2019; Montibus et al., 2023; Ragheb et al., 2020; Saotome et al., 2024), meaning that this could either be a primary function of NuRD or a downstream consequence and/or cell adaptation of NuRD deletion/depletion. For these reasons we have employed a degron system, which allows us to acutely deplete CHD4 protein in mouse ES cells and assess the consequences to chromatin and gene expression within minutes to hours of protein depletion, long before the cells begin to exhibit cell cycle defects.

## Results

### CHD4 has an immediate and widespread impact on chromatin accessibility

NuRD in mouse ES cells can exist in various forms and it can contain different paralogs of key components such as MBD2 or MBD3, GATAD2A or GATAD2B, HDAC1 or HDAC2, and MTA1, 2 and/or MTA3 (Reid et al., 2023). In embryonic stem cells NuRD has only one chromatin remodelling subunit: CHD4. We therefore created ES cell lines in which the endogenous *Chd4* alleles were homozygously tagged with a mini-Auxin inducible degron (mAID) (Nishimura et al., 2009). CHD4-mAID was largely depleted from both the nucleoplasm and chromatin within 60 minutes of Auxin addition (Fig. 1A). CHD4 depleted cells showed a normal cell cycle profile until 24 hours of depletion, when they began to undergo cell cycle arrest at the G1/S checkpoint (Fig. 1B). By 30 hours CHD4-depleted cells were undergoing apoptosis, which increased through 48 hours. We therefore focused our analyses on the first few hours after CHD4 depletion, well before cells started to undergo cell cycle arrest.

**Figure 1.**
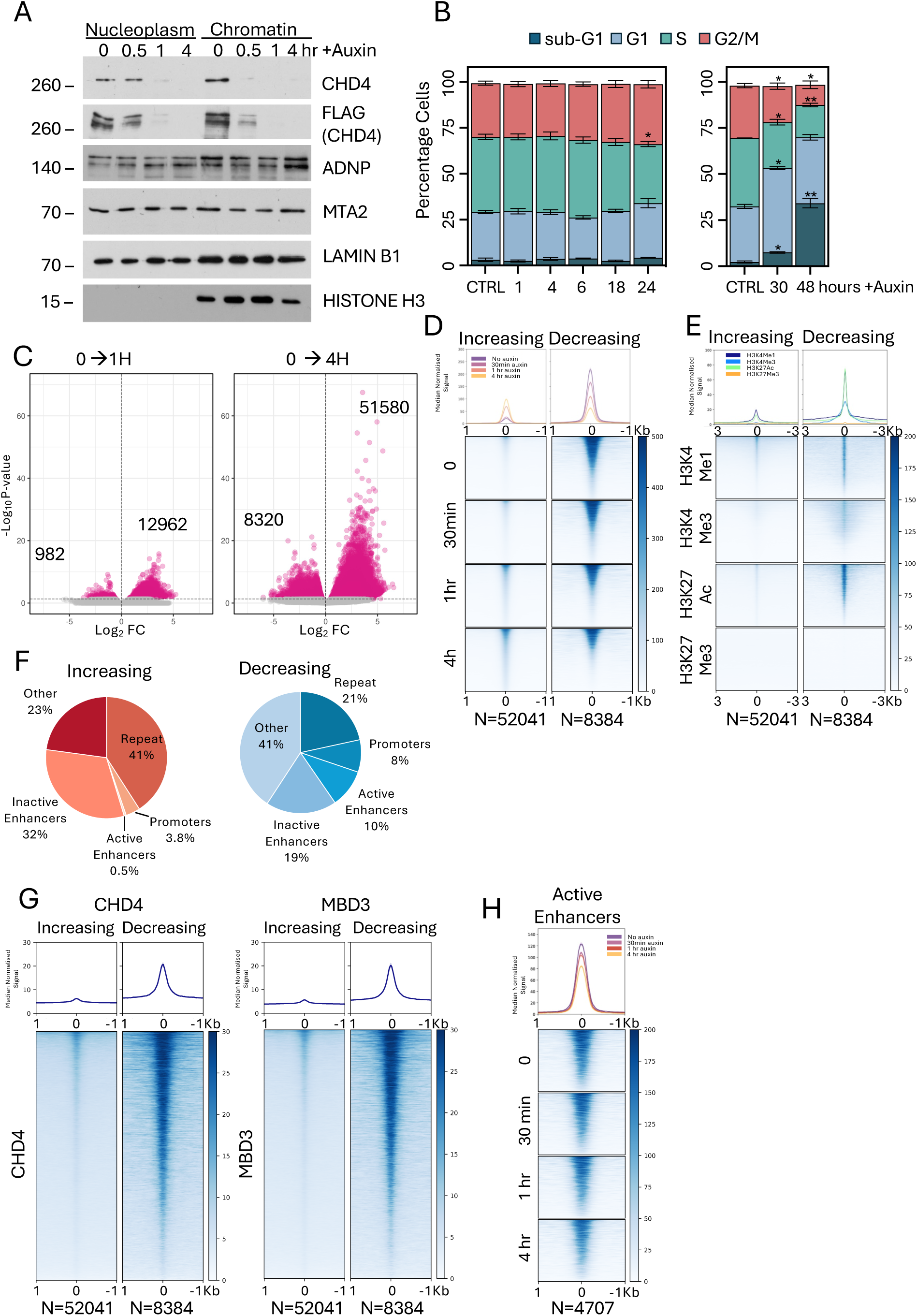
The impact of CHD4 depletion on chromatin accessibility. **A**. Western blots of nuclear soluble (Nucleoplasm) and chromatin fractions across CHD4 depletion probed with antibodies directed against indicated proteins. Times in hours of Auxin addition are indicated across the top. Lamin B1 and Histone H3 act as loading controls. **B**. Cell cycle analysis across CHD4 depletion time course. Hours post Auxin addition are indicated at the bottom, “CTRL” indicates DMSO control. Data represent an average of three replicates. Asterisks indicate significant differences from CTRL using a mixed-effects model with Dunnett’s multiple comparisons correction. *p < 0.05; **p < 0.01. **C**. Volcano plots of differentially accessible ATAC-seq peaks between 0 and 1 hour of Auxin addition (left) or 0 and 4 hours of Auxin addition (right). Magenta spots indicate statistically significant differences (FDR > 0.05). Numbers of significant peaks decreased or increased are indicated on the plots. **D**. Heatmaps of ATAC-seq signal for all regions displaying increased accessibility (N=52041) or decreased accessibility (N=8384) at any time across the CHD4 depletion time course are displayed for each time point. **E**. Heatmaps of CutCRun data for indicated histone modifications at sites increasing or decreasing in accessibility (as in panel **D**) (H3K27Ac and H3K4Me1 from this study; H3K27Me3 and H3K4Me3 taken from (Lando et al., 2024)) at indicated times after CHD4 depletion. **F**. Percentages of sites increasing in accessibility (top, blue) or decreasing in accessibility upon CHD4 depletion (bottom, red) which localise to indicated genomic features. Active enhancers are defined as having H3K4Me1 and K3K27Ac but not K3K4Me3, and inactive enhancers as having H3K4Me1 but not H3K4Me3 or H3K27Ac. **G**. Heatmaps of CHD4 and MBD3 CutCRun data at upDARs and downDARs in 2iL conditions. **H**. Heatmaps of ATAC-seq signal at active enhancers (N=4707) across the CHD4 depletion time course. Median curves in graphs in **D**, **E**, **G** and **H** are plotted with standard error of the mean in lighter shading.

CHD4 depletion had an immediate and widespread impact on chromatin accessibility, as measured by calibrated ATAC-seq. After 60 minutes of CHD4 depletion there were more than thirteen thousand sites showing a significant change in accessibility, which increased to over fifty thousand sites showing increased accessibility and eight thousand showing decreased accessibility by 4 hours (Fig. 1C). These data are consistent with previous reports of overall increased accessibility upon CHD4/dMi2 depletion by shRNA in mouse ES cells or Drosophila S2 cells (Dieuleveult et al., 2016; Moshkin et al., 2012).

Differentially accessible regions increasing in accessibility upon CHD4 depletion exhibited low accessibility and low overall enrichment for specific histone modifications associated with active chromatin in undepleted cells (H3K27Ac, H3K4Me1, H3K4Me3), but not H3K27Me3 (Fig. 1D,E). As a class, these sites predominantly mapped to inactive enhancers and repetitive elements (Fig. 1F). They had very low average enrichment for CHD4 or MBD3 (Fig. 1G) and would likely not pass the threshold to be counted as “Peaks” in many ChIP-seq or CutCRun datasets. Low enrichment for active chromatin marks and NuRD components as well as low but detectable accessibility indicates that these sites predominantly represent silent and cryptic regulatory sequences. Upon CHD4 loss accessibility at these sites increased two-to three-fold at 60 and 240 minutes of depletion (Fig. 1D) indicating that the low level CHD4 enrichment at these sites may nevertheless be functionally relevant.

Sites showing decreased accessibility without CHD4 were highly accessible in undepleted cells, enriched for marks of active chromatin such as H3K4Me1 and H3K27Ac, and to a lesser extent H3K4Me3 (Fig. 1D,E), and often overlapped with known enhancers and promoters (Fig.1F). These sites also showed high enrichment for CHD4 and MBD3 in the undepleted state (Fig. 1G). Upon CHD4 depletion accessibility at these sites was reduced by less than two-fold on average (Fig. 1D). We and others have previously shown that NuRD acts at active enhancers (Basu et al., 2023; Bornelöv et al., 2018; Dieuleveult et al., 2016) and, consistent with these findings, CHD4 depletion caused an average decrease in accessibility at active enhancers (Fig. 1H). We therefore conclude that CHD4 activity maintains closed chromatin generally at inactive regulatory sequences but also contributes to the maintenance of highly accessible chromatin at active sites.

### CHD4 activity maintains expression of active genes while reducing transcriptional noise

Significant changes in gene expression were first detected in nascent and total mRNA 1-2 hours after CHD4 depletion (Fig. 2A,B). Loss of CHD4 resulted in an approximately 2:1 ratio of increased to decreased gene expression from 1 hour onwards (Fig. 2C), consistent with CHD4 activity able to both facilitate and limit transcription, with most genes changing by two-fold or less (Fig. 2D). GO terms associated with activated genes indicate various tissue-specific functions, consistent with the general noise reduction activity described for NuRD (Burgold et al., 2019; Montibus et al., 2023; Ragheb et al., 2020). Downregulated genes, in contrast, are associated with cellular maintenance and early development, consistent with these genes being normally highly expressed in ES cells (Fig. 2E).

**Figure 2.**
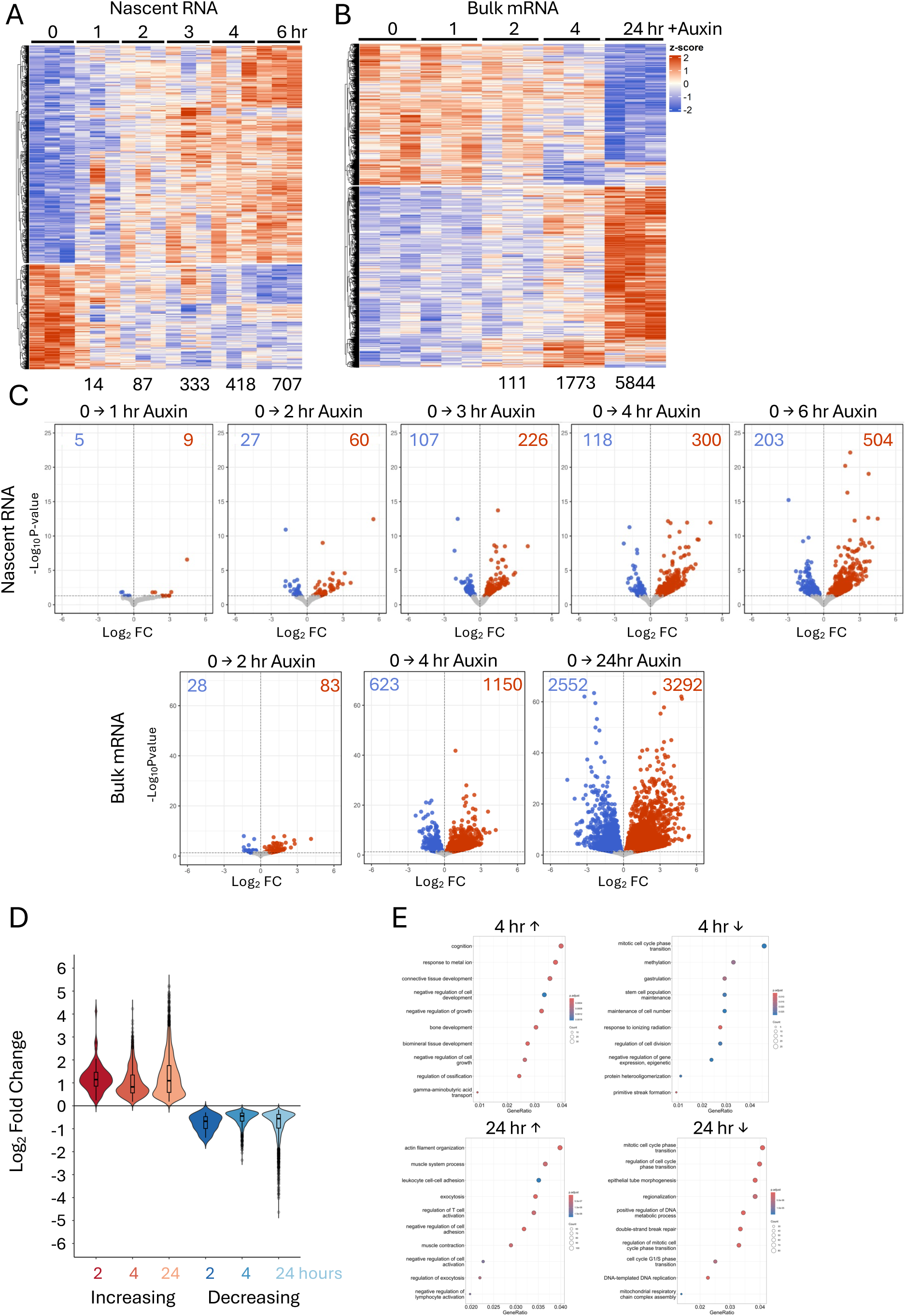
CHD4 acutely regulates gene expression. **A and B.** Heatmaps from nascent RNA-seq (A) or bulk RNA-seq (B) of genes showing significant (P_adj_ <0.05) changes in expression at any point during the CHD4 depletion time course. Heatmaps displays z-scores, meaning expression for each gene has been centred and scaled across the entire time course. **C**. Volcano plots showing significant gene expression changes at indicated time points in nascent RNAseq (top) and bulk RNAseq (bottom). Genes increasing upon CHD4 depletion are shown in red, and those decreasing are shown in blue. The number of significantly misexpressed genes at each time point is indicated in the figure. **D**. Violin plots showing the average Log_2_FoldChange of significant upregulated (red) and downregulated (blue) genes during the CHD4 depletion timecourse. **E**. Gene Ontology (GO) enrichment analysis of genes increased or decreased after four hours or 24 hours of CHD4 depletion. The top 10 biological processes are shown for each category, based on smallest adjusted p-value.

To assess whether the observed changes in chromatin accessibility were linked to changes in gene expression, we plotted the distance between differentially accessible regions and the annotated TSS of genes found to be misexpressed up to 4 hours after CHD4 depletion. Sites increasing in accessibility upon CHD4 depletion tended to be located far from genes repressed by CHD4 (Fig. 3A). This makes it unlikely that CHD4 directly silences gene expression by maintaining condensed chromatin at inactive promoters widely, although it could be acting to prevent activation of distal enhancers. In contrast, a notable number of sites losing accessibility are located within 1Kb of the TSS of misregulated genes (Fig. 3A). This indicates that CHD4 acts at some highly active promoters to maintain both chromatin accessibility and transcriptional fidelity.

**Figure 3.**
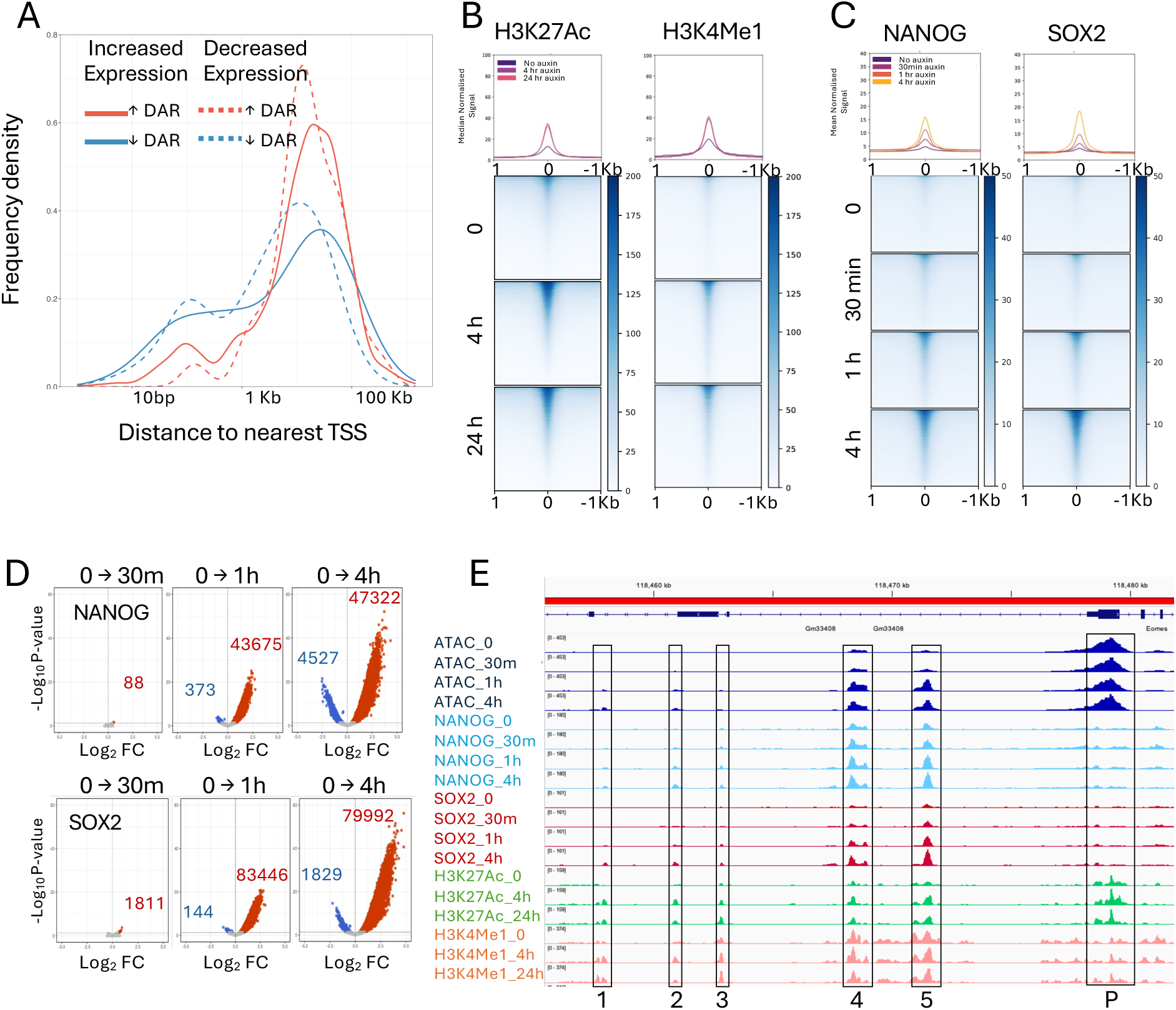
Chromatin opening upon CHD4 depletion. **A**. Frequency density distribution of the distance of increasing Differentially Accessible Regions (“↑ DARs,” red lines) and decreasing regions (“↓ DAR,” blue lines) to the TSS of genes showing increased (solid lines) or decreased (dotted lines) expression within four hours of CHD4 depletion. **B**. Heatmaps of CutCTag signal for H3K27Ac, and H3K4Me1 at sites increasing in accessibility (N = 52041) at indicated times of CHD4 depletion. **C**. Heatmaps of NANOG and SOX2 CutCRun signal at increasing accessibility sites at indicated times of CHD4 depletion. Median curves in **B** and **C** are plotted with standard error of the mean in lighter shading. **D**. Pairwise comparisons of called peaks of binding for NANOG (top) and SOX2 (bottom) between undepleted cells (0h) and 30 minutes, 1 hour or 4 hours of CHD4 depletion. Significantly changed (FDR>0.05) binding sites are shown in blue when log_2_FC>0 and red when log_2_FC<0. **E**. IGV screenshot of the upstream region of the mouse *Eomes* locus displaying ATAC-seq, CutCRun and CutCTag data as indicated at left. Boxed regions labelled 1-5 are CHD4 condensed sites, while the box labelled P corresponds to the *Eomes* promoter.

If loss of CHD4 resulted in activation of normally silent or cryptic enhancers, we would expect that they should show increases in enhancer-associated histone modifications and binding of transcription factors (TFs). As expected, CutCTag for H3K4Me1 and H3K27Ac showed increasing enrichment at CHD4-condensed sites 4 hours after CHD4 depletion, which did not increase further by 24 hours of depletion (Fig. 3B). Although these sites were largely unbound by NANOG or SOX2 in the presence of CHD4, within 1 hour of CHD4 depletion, they became extensively bound by both transcription factors (Fig. 3C). Consistently, thousands of new NANOG and SOX2 peaks were detected across the genome after CHD4 depletion (Fig. 3D). For example, the 25 kb region upstream of the *Eomes* gene contains multiple CHD4-condensed sites (Fig. 3E). Those labelled 1, 2 and 3 show very little, if any, accessibility or TF binding, and very low H3K27 acetylation and H3K4 monomethylation in undepleted cells. All three of these show a gain in accessibility, TF binding and both histone modifications upon CHD4 depletion. Those sites labelled 4 and 5 both show some accessibility, TF binding and enrichment for active histone modifications in undepleted cells. Nevertheless, these sites also show an increase in all of these features upon CHD4 depletion. In contrast, the nearby *Eomes* promoter (“P”) shows little or no change in accessibility, TF binding or enrichment for H3K27Ac or H3K4Me1 across the CHD4 depletion time course.

We propose that low-level association of CHD4 and NuRD across euchromatin restricts chromatin accessibility, preventing transcription factors from binding to consensus sequence motifs. It is also possible, however, that accessibility increases are a consequence of more stable transcription factor binding. Failure to maintain this level of chromatin inaccessibility results in activation of inactive and/or cryptic enhancers, which can stimulate inappropriate gene expression.

### CHD4 and SALL4 maintain chromatin inaccessibility largely independently

The SALL4 protein is a very abundant transcription factor in mouse ES cells, which has long been known to interact with the NuRD complex (Kloet et al., 2015; Lauberth and Rauchman, 2006; Lu et al., 2009; Miller et al., 2016). SALL4 preferentially binds to A/T-rich DNA genome-wide, where it is proposed to exert a general repression function through NuRD recruitment (Kong et al., 2021; Pantier et al., 2021; Ru et al., 2022; Watson et al., 2023). If this model is correct, we would expect some proportion of the CHD4 condensed sites to be dependent upon both CHD4 and SALL4 to remain inaccessible. To test this model, we assessed chromatin accessibility after dTAG-mediated SALL4 depletion. SALL4 is partially redundant with SALL1 in ES cells, but only SALL4 is required for early mammalian development (Miller et al., 2016; Nishinakamura et al., 2001). We therefore created *Sall1^-/-^Sall4^FKBP/-^* ES cells in which one endogenous *Sall4* allele was mutated and the remaining allele was targeted to express a SALL4-FKBP fusion protein.

Acute depletion of SALL4-FKBP resulted in rapid and extensive changes in chromatin accessibility as measured using ATAC-seq. Over 18,000 sites showed increased accessibility within 1 hour of SALL4 depletion in *Sall1^(-/-)^* ES cells (Fig. 4A,B). These sites showed moderate enrichment for SALL4 in undepleted cells, though less than that seen at active enhancers (Fig. 4C). When we compared these sites to those showing CHD4-dependent chromatin condensation, we find that only 4559 sites of the more than 52,000 CHD4 -dependent sites also show dependency upon SALL4 to maintain chromatin inaccessibility (Fig. 4D-F). This means that only 17.2% of SALL4-dependent sites also rely on CHD4 to prevent chromatin opening. Moreover, over 47,000 CHD4-dependent sites show neither SALL4 binding nor SALL4 dependence to remain inaccessible (Fig 4. D-G). Similarly, over 20,000 SALL4-dependent sites show no change in accessibility upon CHD4 depletion, despite showing similar levels of enrichment for NuRD components as those dependent upon both SALL4 and CHD4 (Fig. 4E,H). Consistent with the proposed preference of SALL4 for binding to A/T-rich DNA, SALL4-unique regions showed enrichment for SALL4 binding and a high A/T-content (53.9%) (Fig. 4D). CHD4-unique regions showed little if any SALL4 enrichment and a much lower A/T content (44.2%), while those sites requiring both SALL4 and CHD4 to remain inaccessible showed enrichment for NuRD components as well as SALL4 and displayed intermediate A/T content (48.2%). Together, these data show that both CHD4 and SALL4 broadly restrict chromatin accessibility across the genome, but that their activities overlap at only a relatively small proportion of sites.

**Figure 4.**
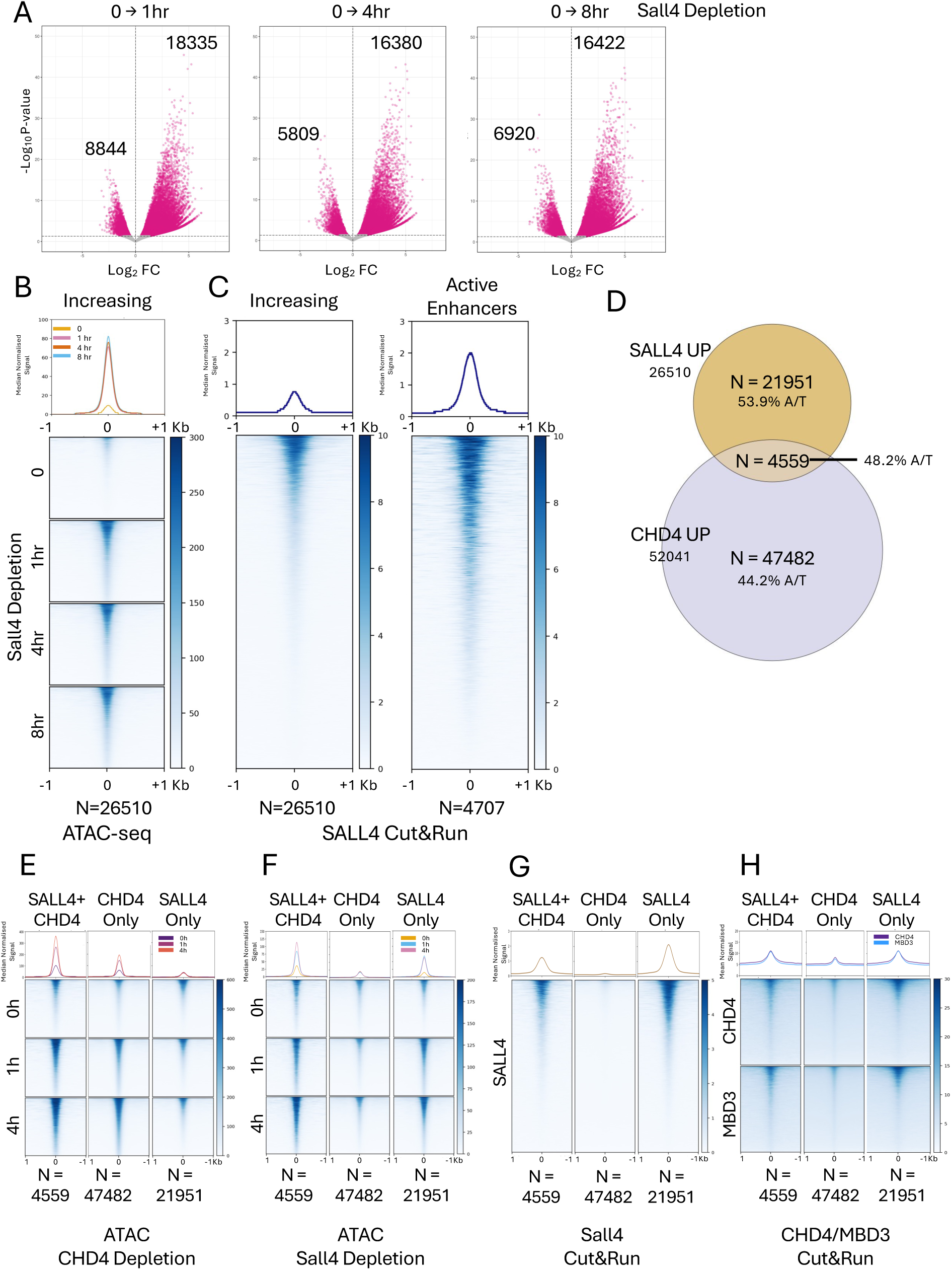
CHD4 and SALL4 both restrict chromatin accessibility. **A**. Volcano plots of differentially accessible ATAC-seq peaks when comparing 1 hour, 4 hours or 24 hours of SALL4 depletion with those seen in undepleted cells (0). Magenta spots indicate statistically significant differences (FDR > 0.05). Numbers of significant peaks decreased or increased are indicated on the plots. **B**. Heatmaps of ATAC-seq signal for all regions displaying increased accessibility (N=26510) across the SALL4 depletion time course are displayed for each time point. **C**. Heatmap of SALL4 CutCRun signal in undepleted ES cells (taken from (Ru et al., 2022)) at all regions displaying increased accessibility across the SALL4 depletion time course (left, N=26510) or at active enhancers (right, N=4339). **D**. Overlap of sites showing increased accessibility upon SALL4 depletion with those increasing upon CHD4 depletion (upDARs). The % A/T base composition of the different categories of sites is indicated. **E, F**. Heatmaps of ATAC-seq signal at sites increasing upon either SALL4 or CHD4 depletion (SALL4+CHD4), sites increasing upon CHD4 depletion but not upon SALL4 depletion (CHD4 Only), or sites increasing upon SALL4 depletion but not upon CHD4 depletion (SALL4 Only) plotted at indicated time points of CHD4 depletion (**E**) and SALL4 depletion (**F**). **G, H**. CutCRun signal for SALL4 (**G**) or for CHD4 and MBD3 (**H**) in undepleted ES cells at the three different classes of sites. Median curves in **B, C,** and **E-H** are plotted with standard error of the mean in lighter shading.

### NuRD limits transcription factor binding to chromatin

Sites decreasing in accessibility upon CHD4 depletion showed an increase in enrichment for both NANOG and SOX2 after 30 minutes, even though these sites were, on average, losing accessibility and losing enrichment for the active chromatin marks H3K4Me1 and H3K27Ac at that time (Fig. 5A,B). Active enhancers similarly showed an increase in NANOG and SOX2 enrichment from 30 minutes of CHD4 depletion (Fig. 5C). An example locus is shown in Figure 5D, where a cluster of enhancers located 50-70Kb downstream of the *Klf4* gene contains several high accessibility sites. Those located at 68, 57 and 55 Kb downstream of *Klf4* (labelled 68, 57 and 55, respectively in Figure 5D) all show a loss of ATAC-seq signal but a gain in NANOG and, to a lesser extent, SOX2 enrichment between 0 and 30 minutes of CHD4 depletion. NANOG enrichment then decreases again at 1 and 4 hours while Sox2 enrichment remains relatively constant, even though accessibility continues to decrease.

**Figure 5.**
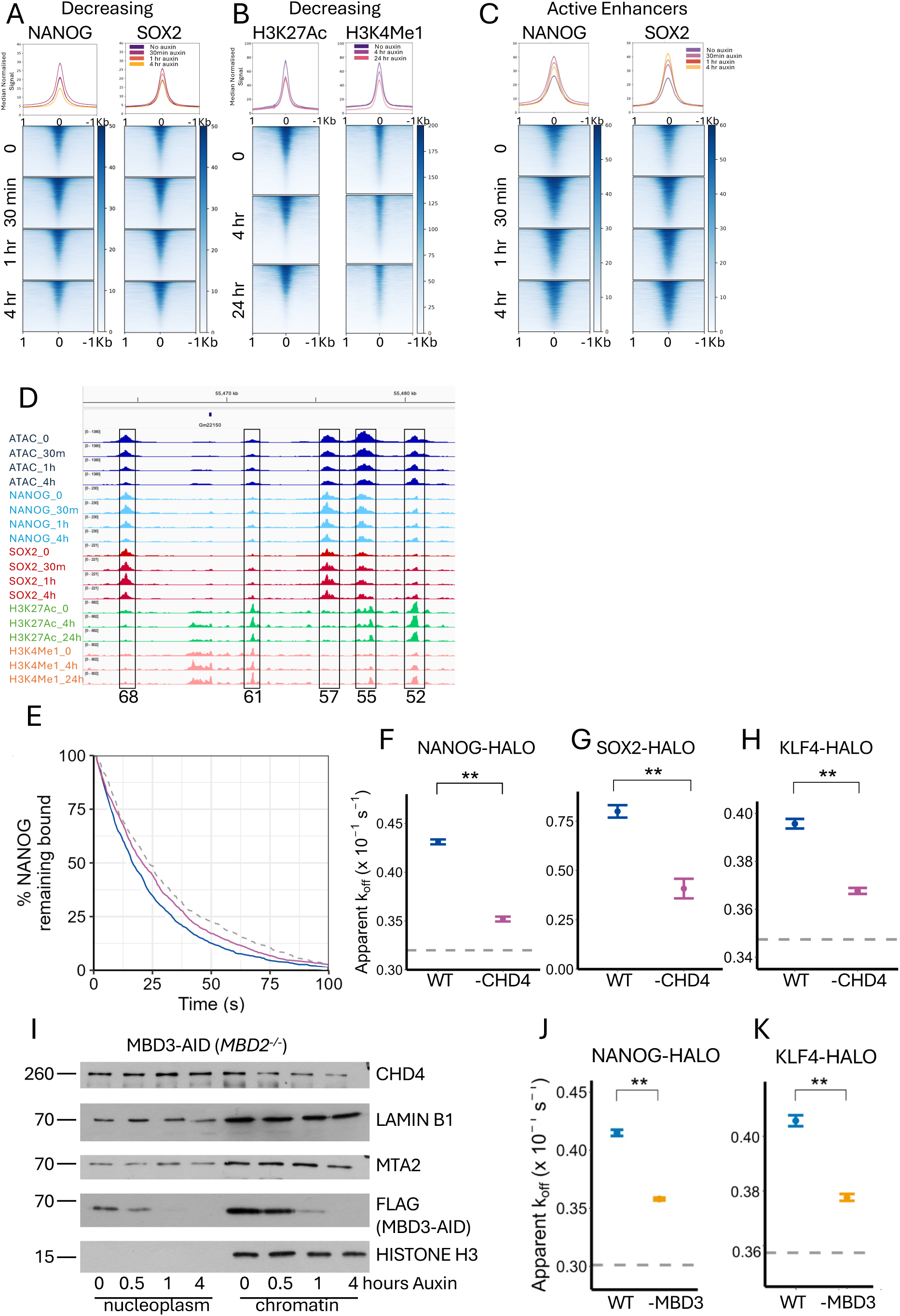
NuRD regulates NANOG and SOX2 binding to active sites. **A**. Heatmaps of NANOG and SOX2 CutCRun signal at indicated times of CHD4 depletion across sites decreasing in accessibility (N = 8384). **B**. Heatmaps of H3K27Ac and K3H4Me1 CutCTag signal at indicated times of CHD4 depletion across decreasing accessibility sites. **C**. Heatmaps of CutCRun signal for NANOG and SOX2 across active enhancers (N = 4707) at indicated times of CHD4 depletion. Median curves in **A-C** are plotted with standard error of the mean in lighter shading.**D.** IGV screenshot of the enhancer cluster downstream of the *Klf4* gene displaying ATAC-seq, CutCRun and CutCTag data as indicated at the left. Boxed regions are labelled with the distance in Kb from the annotated *Klf4* transcription start site. **E**. Fluorescence survival curves of chromatin-bound NANOG-HALO molecules in 2iL (blue line) or after 1 hour of CHD4 depletion (purple line). The grey dotted line represents the fluorescence survival curve for molecules in a fixed-cell control imaged under identical conditions. **F**. Apparent dissociation rates (k_off_) of chromatin-bound NANOG molecules calculated through fitting a single exponential decay model to the survival curves in **panel E**. Error bars represent 95% confidence intervals for each fit applied to data taken from three independent experiments. The horizontal dashed line represents the upper 95% confidence limit for a fixed-cell control. ** indicates that 99% confidence intervals do not overlap, i.e. p < 0.01. **G.H**. As in **F** but for SOX2 (**G**) and KLF4 (**H**). The fixed cell control was not imaged in the SOX2 experiments in panel **F**. **I**. Western blots of nuclear soluble (Nucleoplasm) and chromatin fractions across MBD3 depletion probed with antibodies directed against the indicated proteins. Times in hours of Auxin addition are indicated across the top. Lamin B1 and Histone H3 act as loading controls. **J, K.** Apparent dissociation rates (k_off_) for NANOG-HALO (**I**) and KLF4-HALO (**J**) before and after 60 minutes of MBD3 depletion. For calculation of k_off_ the trajectories were pooled from 4 replicates of each time point obtained over 2 days.

The simultaneous decrease in chromatin accessibility and increase in transcription factor enrichment after 30 minutes of CHD4 depletion could indicate that CHD4/NuRD plays a direct role in controlling transcription factor binding to chromatin. To independently assess the impact CHD4 has on transcription factor binding, we used single molecule imaging in live ES cells to directly measure transcription factor binding kinetics upon CHD4 depletion. A HALO-Tag cassette was knocked-in to the endogenous *Nanog*, *Klf4* or *Sox2* loci in the CHD4-mAID ES cell line to create strains expressing the different C-terminal protein fusions. Single HALO-tagged protein molecules were then labelled with a photoactivatable dye (PA-JF646, (Grimm et al., 2016)) and tracked at two distinct temporal regimes where we collected images either every 20 ms or 500 ms, using double-helix point spread function (DH PSF) microscopy as they moved within a 4-μm slice of the nucleus. Recording at a 20 ms time resolution allows the segmentation of trajectories into freely diffusing and chromatin-bound states, and this data can be used to extract the chromatin binding kinetics of proteins and complexes (Basu et al., 2023). Here, we used these data to determine residence times – how long the molecules remain bound to chromatin. The proportion of molecules that remain bound to chromatin after increasing lengths of time was then plotted before and after CHD4 depletion (for 1 hour) and the results were fitted with a single exponential decay to determine the apparent dissociation rate, k_off_ (Fig. 5E). NANOG-HALO, SOX2-HALO and KLF4-HALO all showed a pronounced decrease in apparent k_off_ after CHD4 depletion (Fig. 5F-H), consistent with CHD4 actively limiting transcription factor residence times on chromatin.

To determine whether it is CHD4 by itself or the intact NuRD complex that can remodel transcription factor binding, we similarly imaged NANOG-HALO and KLF4-HALO in MBD3-depletable ES cells after disruption of the NuRD complex (Fig. 5I). These cells were also null for *Mbd2*, which encodes a protein displaying partial functional redundancy with MBD3 but is dispensable in ES cells (Hendrich et al., 2001). As CHD4 is largely responsible for the binding of NuRD to its target enhancers, removal of MBD3 enables disassembly of NuRD, but does not prevent CHD4 binding to chromatin (Basu et al., 2023; Bornelöv et al., 2018; Zhang et al., 2016). The nucleosome remodelling activity of CHD4 is, however, greatly reduced outside of NuRD (Bornelöv et al., 2018). We therefore reasoned that if it is intact NuRD that is required (as opposed to just CHD4), depletion of MBD3 should also limit TF residence times. Indeed, we detected a significant decrease in the apparent k_off_ of stably bound NANOG-Halo and KLF4-Halo after 1 hour of MBD3 depletion (Fig. 5J,K). We therefore conclude that intact NuRD (and not CHD4 by itself) actively limits transcription factor residence times on chromatin, while at the same time contributing to the maintenance of chromatin accessibility at active regulatory regions.

### NuRD activity influences active and inactive regulatory elements differently

We next asked how CHD4 activity could have different impacts on accessibility at different kinds of sequences. Information about the structure of accessible regions can be obtained from ATAC-seq data by quantitating recovered fragments of all sizes and determining the frequency and location of recovered Tn5 integration sites using VPlots, (Henikoff et al., 2011; Schep et al., 2015; Serizay and Ahringer, 2021)(Supplementary Figure 2). Applying this analysis at CHD4 -ondensed sites shows an increase in both Tn5 integrations and in reads corresponding to the nucleosome-free region (NFR) after 30-minutes of CHD4 depletion (Fig. 6A,B). After 60 minutes, when CHD4 is almost completely depleted, there is an increase in accessibility generally across the entire site, both of short reads within the NFR but also longer reads (200-300bp) extending from the NFR outwards (Fig. 6B). We conclude that NuRD is acting to restrict accessibility of both the NFR and the flanking nucleosomal DNA (see Supplementary Figure 2) at these sites, consistent with the demonstrated role of CHD4/NuRD in maintaining the density of intact nucleosomes (Bornelöv et al., 2018; Dieuleveult et al., 2016; Moshkin et al., 2012).

**Figure 6.**
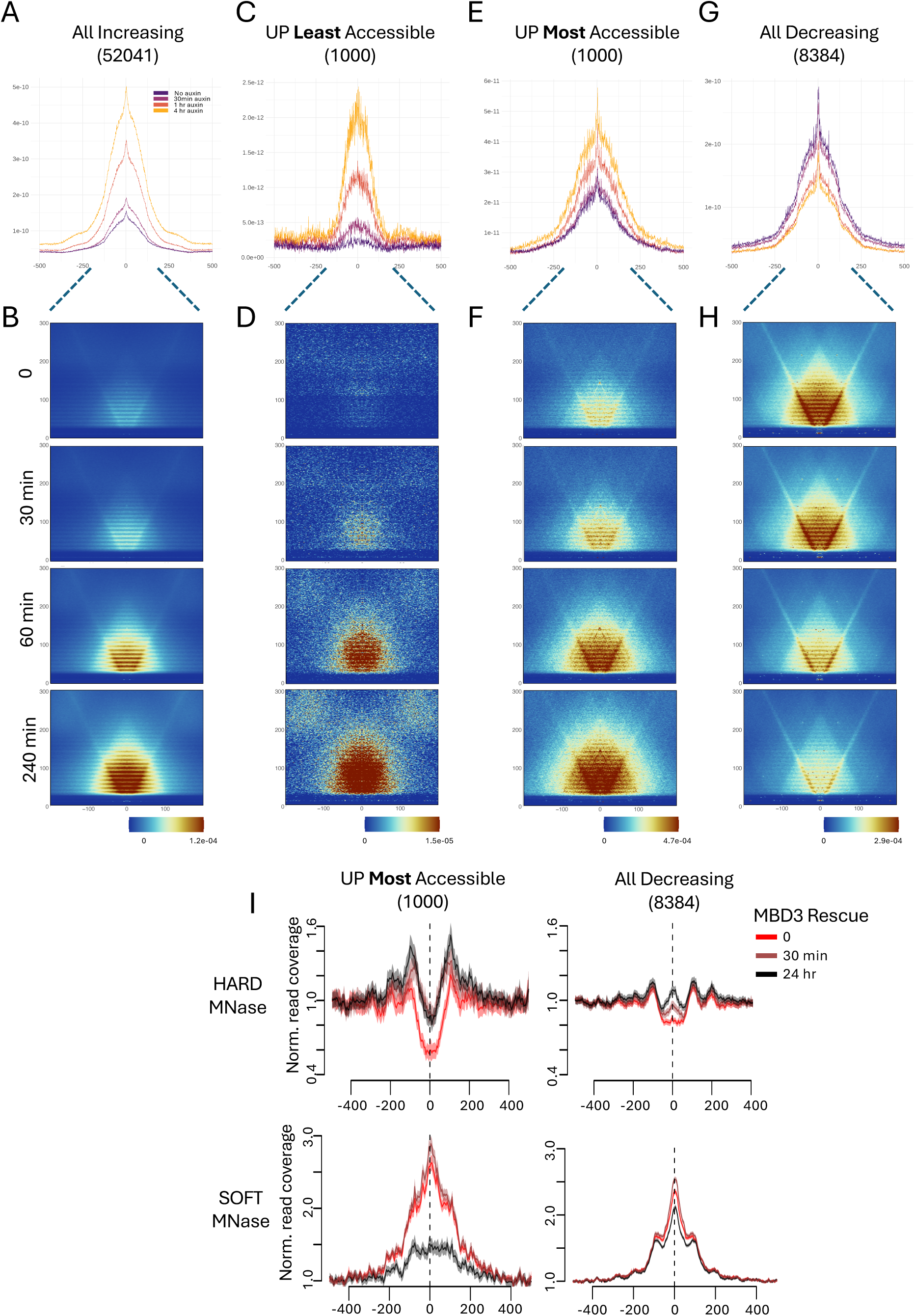
CHD4 controls accessibility differently at different classes of sites. **A**, **C, E, G** Tn5 integration frequency was determined from ATAC-seq data and plotted across indicated sites across the CHD4 depletion time course. The number of sites is shown in parentheses. **B, D, F, H** Vplots across sites indicated above (corresponding to the Tn5 integration plots) at indicated times of CHD4 depletion. See **Supplementary Figure 2** for a Vplot schematic. **I.** MNase-seq data collected 0 (red), 30 minutes (dark red) or 24 hours (black) after of NuRD reformation in *Mbd3^-/-^* ES cells are plotted across indicated sites. Top graphs show results from “Hard” MNase treatment while bottom graphs show “Soft” MNase treatment (see text). The y-axis shows normalised read coverages, while the X-axis shows distance in base pairs from the centre of the feature. Curves show mean and standard error from 3 biological replicates.

The example of the *Eomes* locus (Fig. 3F) illustrates that some CHD4-condensed sites show a degree of accessibility in the presence of CHD4, while others show little if any accessibility. To better understand how CHD4 prevents chromatin opening at these two different kinds of sites, we constructed Tn5 integration plots and Vplots from the 1000 CHD4-condensed sites showing the least accessibility in undepleted conditions and from the 1000 showing the most accessibility (Fig. 6C-F). Sites showing little or no accessibility in undepleted cells initially show very few reads less than 100bp. The V-plots also show reads of 200-300 bp distributed across the regions, again indicative of generally inaccessible chromatin. Loss of CHD4 results in the formation and progressive expansion of an NFR, as well as an increase in longer reads with one end located within the NFR (Fig. 6C,D). The Tn5 integration plot does not show a general widening of the NFR over time, indicating that CHD4 function is largely focussed on maintaining nucleosome density within a 100-200bp region (Fig. 6C). Sites at which CHD4 activity restricts accessibility at already accessible sites showed a general increase in both small and longer reads as CHD4 was depleted, and a broadening of the existing NFR (Fig. 6 E,F). Together these data are consistent with CHD4/NuRD maintaining chromatin compaction at inactive or low activity enhancers by maintaining nucleosome density.

The sites at which CHD4 activity maintains accessibility displayed a prominent NFR as well as high density of longer reads in undepleted cells (Fig. 6G,H). CHD4 depletion induced a general decrease in Tn5 integrations and VPlot signal intensity across these sites, consistent with CHD4 acting to generally maintain accessibility.

We next asked why CHD4 promoted accessibility at highly accessible active sites (e.g. Fig. 6G,H) but reduced accessibility at slightly less accessible, inactive sites (e.g. Fig. 6E,F). We took advantage of ES cells in which a tamoxifen-inducible MBD3b is expressed in an otherwise *Mbd3*-null ES cell line, allowing us to restore NuRD activity to cells upon tamoxifen addition (Bornelöv et al., 2018; Reynolds et al., 2012). We had previously subjected these cells to micrococcal nuclease (MNase) sequencing at different time points after MBD3 induction to show that restoration of NuRD activity caused increased density of intact nucleosomes across sites of active transcription (Bornelöv et al., 2018). Unlike ATAC-seq, sequencing of MNase-treated DNA results in increased reads at sites protected from digestion by bound proteins, i.e. less accessible regions, and a decrease in signal at locations where the DNA is easily accessible and therefore digested by the MNase. In that study, we used an MNase concentration at which DNA associated with intact nucleosomes is protected from MNase digestion. Lower MNase concentrations, however, will recover DNA protected by other chromatin proteins as well as by partial (fragile) nucleosomes (Kubik et al., 2015; Xi et al., 2011).

We therefore subjected cells undergoing the time course of NuRD reformation to lower MNase concentrations prior to sequencing. Traditional MNase sequencing shows that NuRD increases the density of intact nucleosomes at both classes of accessible sites, as indicated by an increase in signal from the red line (NuRD-deficient) to the black line (24h NuRD restored) in the top panels of Figure 6I. This effect is more pronounced at the most accessible inactive sites (“UP Most Accessible”, black line, Fig. 6I), occurring across a broader area than at fully active sites (“All Decreasing”, black line). The lower MNase concentration, by contrast (Fig. 6I, lower panels) produces a peak of MNase resistance in the absence of NuRD activity at both classes of sites (red lines), indicating an accumulation of structures which are digested by the higher MNase concentration, such as fragile nucleosomes. Restoration of NuRD activity clears the majority of this signal at the most accessible inactive sites, but only moderately reduces their abundance at fully active sites (Fig. 6I, lower panels). These data indicate that at highly active regions, NuRD acts to limit the abundance of fragile nucleosomes and other proteins, while marginally increasing the density of intact nucleosomes, which has an overall result of maintaining these sites in a fully active state. It does not completely remove the fragile nucleosomes from these sites, which are presumably being created by the activity of other chromatin remodellers (Klein et al., 2023; Nocente et al., 2024). In contrast, at accessible but inactive sites it prevents the accumulation of fragile nucleosomes which would otherwise cause further opening and enable activation of inactive regulatory elements. This means that the activity of CHD4 does not differ at the two classes of sites, but it is the presence of other remodellers producing fragile nucleosomes which dictates whether CHD4 activity acts to maintain or prevent accessibility (Fig. 7).

**Figure 7.**
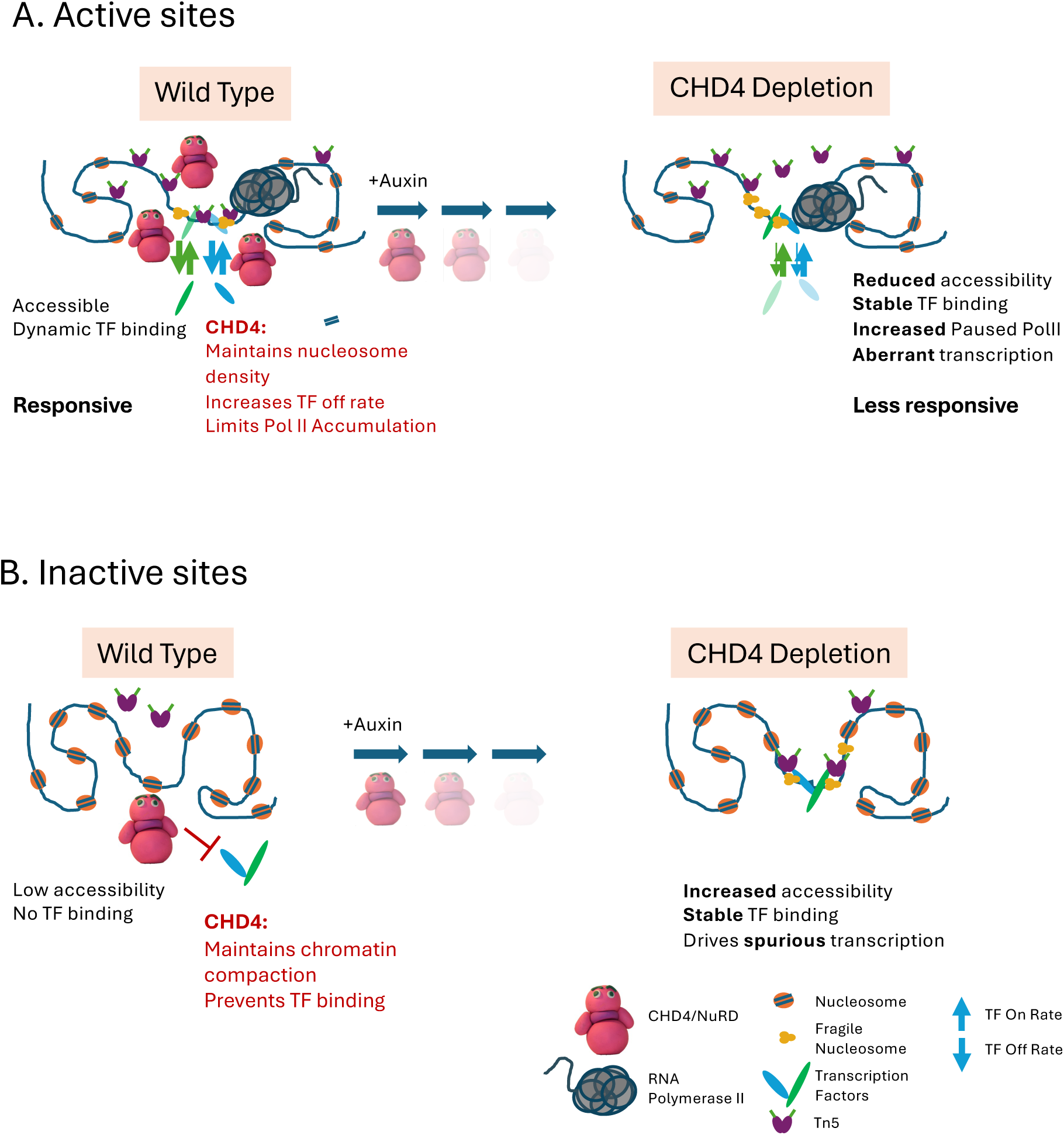
Model of CHD4 function. **A. Highly accessible sites.** In undepleted cells these sites are extensively bound by CHD4/NuRD, where it acts to promote the off rate of transcription factors to promote accessibility. Tn5 is able to access the central NFR but also can integrate into the flanking nucleosomal DNA. After CHD4 depletion, the on rate for TFs does not change, but the off rate is now much reduced, resulting in increased TF binding. The sites become less accessible to Tn5, such that although it can still access the hypersensitive site within the NFR, there are fewer integrations extending outwards. These regulatory regions cannot quickly respond to receipt of external signals. **B**. Model of CHD4 function at inaccessible, silent enhancers. In undepleted conditions, there is low CHD4 enrichment at these sites. Here, CHD4 acts to prevent binding of transcription factors and maintain low accessibility, such that Tn5 cannot frequently access the DNA. After CHD4 depletion, the locus becomes more accessible, and transcription factors can stably bind. This leads to spurious activation of distal promoters and an increase in transcriptional noise.

Together, these data show that CHD4 shows two modes of binding on chromatin: in addition to the high enrichment seen at active enhancers and promoters, it also shows low-level binding at silent and/or cryptic regulatory elements across euchromatin. At the former class of sites, it acts to limit the residence times of chromatin-associated proteins to maintain highly accessible chromatin. This maintenance activity of CHD4/NuRD is necessary for the full activity of these elements to direct appropriate transcription of target genes. At the latter class of sites we propose it maintains chromatin in an inaccessible state, preventing accumulation of fragile nucleosomes and preventing transcription factors from recognising and stably binding their cognate motifs and thereby prevents spurious activation of these elements.

## Discussion

Assessing CHD4 function by traditional knockout experiments is difficult as ES cells without CHD4 undergo cell cycle arrest and begin to apoptose after 24 hours (Fig. 1B). Even with siRNA or inducible deletion, cells in a population will lose CHD4 at different times and, in the case of siRNA, to different extents, making primary function difficult to ascertain. Genetic deletion in mouse ES cells has been used extensively to define protein function in cells and in intact mice. In such an experiment the genetic lesion will often be induced in a single cell, and a clone of cells will be grown out from that single cell. While this is an extremely powerful method for defining gene function, there is usually a very large number of cell divisions occurring and considerable time passing between loss of the gene and assessment of phenotype, making it of limited use when studying proteins required for the viability of ES cells. Resulting phenotypes may be due directly to the absence of protein function but may also reflect adaptations made by cells selected to proliferate in the absence of that protein. The advent of inducible depletion methods (summarised in (Wit and Nora, 2023)) has meant we can now infer protein function by assessing the immediate molecular consequences of loss of a given protein activity in the very short term. Combined with genome-wide profiling, this has allowed us to identify both a novel transcription factor remodelling activity for CHD4 and to define how this combines with the protein’s nucleosome remodelling activity to exert different functions at different classes of sites.

CHD4 is an abundant chromatin remodeller with a well-defined function in moving intact nucleosomes. Here we show that CHD4 can also actively limit the residence times of transcription factors on chromatin. The combination of these two activities has different consequences at different kinds of sequences. At highly accessible, active sites (Fig. 7A) CHD4 enrichment is high and it acts to limit the binding of transcription factors and fragile nucleosomes to regulatory regions, allowing for full accessibility and functionality of these regulatory elements. At inactive, less accessible euchromatic sites (Fig. 7B) CHD4 associates more infrequently, but here it’s TF removal activity, combined with maintenance of nucleosome structure, is sufficient to keep accessibility low and to prevent transcription factors from stably binding recognition motifs. This activity is sufficient to prevent the binding of NANOG and SOX2, the latter being capable of binding nucleosomal templates (Dodonova et al., 2020; Zhu et al., 2018). By combining these two activities, CHD4 functions to prevent spurious activation of cryptic and/or silent regulatory regions, thereby reducing transcriptional noise, but also acts to facilitate the activity of highly accessible regulatory regions, allowing cells to accurately and rapidly respond to differentiation cues (Fig. 7).

Many transcription factors recognise and bind to specific sequence motifs in DNA. A transcription factor recognising a 7 base pair sequence should, on average, have more than 150,000 different potential binding sites in a mammalian genome. Yet ChIP-seq and CutCRun studies have found most transcription factors associate with orders of magnitude fewer sites in any one cell type (Lambert et al., 2018; Srivastava and Mahony, 2020). Most transcription factors bind preferentially or exclusively in accessible chromatin, so it is believed that differences in chromatin accessibility, or nucleosome positioning across sites, determines which subset of the potential consensus DNA binding motifs are available for TF binding at any given time or cell type (Workman and Buchman, 1993; Zhu et al., 2018). Here we identify CHD4 as one factor which acts to prevent transcription factors, even so-called Pioneer Factors, from binding thousands of sites in inaccessible chromatin. Consistently with this model, CHD4 was found to prevent binding of the GATA3 transcription factor to inappropriate sites in breast cancer cells undergoing mesenchemal-to-epithelial transition (Saotome et al., 2024). We propose that CHD4’s chromatin remodelling activity keeps the sites inaccessible, so TF binding is not favoured (i.e. k_on_ is low). The ability of CHD4 to remove TFs from chromatin means that even if they are capable of binding to a nucleosomal substrate, CHD4 promotes their dissociation from chromatin so they are quickly removed (Fig. 7B). At highly accessible chromatin, CHD4 similarly limits TF residence times by increasing k_off_, however accessibility is high so the k_on_ is also high. We propose that at these sites CHD4 activity balances the high k_on_ by maintaining a high k_off_, thus facilitating the turnover of bound molecules (Fig. 7A). Without CHD4, k_off_ decreases while k_on_ remains high, resulting in more stable binding of TFs to chromatin. CHD4 and NuRD have been shown to be important for cells to properly respond to differentiation cues in a wide variety of organisms and contexts (Ahringer, 2000; Bracken et al., 2019; Lenz and Brehm, 2023; McDonel et al., 2009; Signolet and Hendrich, 2015) and we propose that by facilitating TF turnover and accessibility at promoters and enhancers, CHD4 and NuRD facilitate the ability of regulatory elements to respond to differentiation cues (Fig. 7).

CHD4’s activity to prevent spurious TF binding and chromatin opening occurs at many sites where enrichment of CHD4 by ChIP or CutCRun is low, but not absent. Despite the low amount of CHD4 detected at these sites, this activity was sufficient to prevent activation of silent and cryptic regulatory regions (Fig. 2). This low-level enrichment of CHD4 at sites where it prevents activation could explain an observation made during Drosophila spermatogenesis, where the CHD4 orthologue dMi2 was found to prevent transcription from cryptic promoters, despite not being detected at these promoters by ChIP-seq assays (Kim et al., 2017). A recent study in *S. cerevisiae* found many transcription factors appear to exert influences over expression of genes without detectable association at that gene and bind many sites where they exert no detectable regulatory function (Mahendrawada et al., 2025). While some have argued that one should focus on high confidence binding targets to assess protein function (Mindel and Barkai, 2025), in our case this would have resulted in us ignoring most of the 50,000 sites at which we detect CHD4-dependent chromatin compaction (Fig. 1C). While we have argued that chromatin changes detected within 30 minutes are likely to be direct consequences of NuRD manipulations (Fig. 6), we do acknowledge that it is possible these are indirect consequences of loss or gain of NuRD activity. We agree that the definition of “targets” can be highly variable between studies, which is why we preferred to assess CHD4 enrichment at subsets of sites where function was identified, irrespective of whether it passed the threshold of being a “peak.”

SALL4 presents another example of a protein exerting function away from its predominant sites of chromatin enrichment: initial ChIP-seq results indicated that SALL4 was predominantly located at enhancers, whereas more recent CutCRun data indicates a wider spectrum of sites bound by SALL4 (Kong et al., 2021; Miller et al., 2016; Pantier et al., 2021; Ru et al., 2022). We find that SALL4 shows lower enrichment at A/T-rich genomic sites than it does at enhancers, but that it nevertheless has a major impact on chromatin compaction at these genomic A/T-rich sites (Fig. 4). While many groups, including ours, have generally assumed protein function would be focussed at ChIP-seq “peaks,” we argue here that focusing on protein enrichment levels on chromatin is not necessarily the best way to identify important sites of protein activity.

Together, our data show that CHD4 exerts functions on chromatin beyond its well-described ability to slide intact nucleosomes along the DNA (Zhang et al., 2016; Zhong et al., 2020). Consistent with this conclusion, a recent paper found that acute depletion of CHD4 resulted in an increase in fragile nucleosomes at enhancers (Nocente et al., 2024), while we and others have found that NuRD acts to limit the amount of paused RNA Polymerase II at active promoters (Bornelöv et al., 2018; Pundhir et al., 2023). It is possible that CHD4 translocates along the DNA within accessible sequences and displaces any bound proteins, such as transcription factors or fragile nucleosomes, that it encounters. Both transcription factors and fragile nucleosomes are less tightly bound to DNA than intact nucleosomes, so it is possible CHD4’s nucleosome remodelling activity results in their displacement, while still sliding intact nucleosomes. CHD4 could also displace transcription factors from preferred binding sites, thus making binding less favourable. Such activity might not be necessary in inaccessible chromatin, but at inactive enhancers showing a small amount of accessibility, infrequent binding and translocation of NuRD across the accessible region could remove any proteins which have managed to bind. Whether this is achieved through a similar ATP-dependent mechanism as is used to slide nucleosomes remains to be determined (Reid et al., 2024; Zhong et al., 2020). Notably, the yeast remodeller INO80 was found to bind differently to fragile nucleosomes vs intact nucleosomes, possibly indicating different mechanisms of remodelling the two different substrates (Wu et al., 2023; Zhang et al., 2023).

A recent study showed that the nucleosome remodelling activity of a different remodeller, SMARCA5, is dictated by the density of nucleosomes on the DNA that it encounters: at high density sites it maintains that density, while at low density sites it slides nucleosomes across the DNA to facilitate TF access (Abdulhay et al., 2023). We find that CHD4 increases density of intact nucleosomes at sites where it prevents or maintains accessibility (Fig. 6I, top panels) and also limits TF residence times at both classes of sites. Therefore, unlike SMARCA5, we see no evidence that the activity of CHD4 differs between condensed and accessible chromatin, but rather the consequences of that activity are different at different kinds of sites (Fig. 7).

The residence times of transcription factors vary but are generally on the order of a few seconds (Lu and Lionnet, 2021). Why might it be important to limit the residence times of transcription factors? We propose that this could fix an enhancer into a specific state, when one important job of enhancers is to be responsive to changes in stimuli. If no change in status quo is required, then after CHD4 promotes eviction of a particular transcription factor from the enhancer it can continually re-bind and exert its influence over that enhancer. Should signals change, the enhancer needs to be able to remove TFs corresponding to the old signal to make space for determinants of the new signal. In this model, CHD4/NuRD maintains this fluidity of transcription factor interactions, ensuring enhancers are rapidly able to respond immediately upon receipt of differentiation cues.

## Methods

### Mouse embryonic stem (ES) cells

Mouse ES cells were cultured in 2i+LIF media on gelatin-coated plates (Montibus et al., 2023; Mulas et al., 2019). All cell lines were genotyped and tested for mycoplasma regularly. The CHD4-mAID ES cell line was made in BC8, an F1 hybrid from a C57Black/6 and Mus castaneus cross (40, XY), obtained from Anne Ferguson-Smith (Cambridge). MBD3-AID ES cells were created in 23AF, a primary ES cell line derived from *Mbd2^-/-^*, *Mbd3^Flox/Flox^* mice (40, XX) in a mixed C57Black/6 and 129/Ola background. Sall4-FKBP cells were created in *Sall1^-/-^ Sall4^+/-^* ES cells (Miller et al., 2016).

Targeting constructs were made using the AID sequence (Nishimura et al., 2009) or mini-AID (Kubota et al., 2013) amplified from an Oct4-AID plasmid; a gift from José Silva (Bates et al., 2021). Targeting plasmids using the FKBP protein (Nabet et al., 2018) were made by amplifying FKBP from pLEX_305-C-dTAG; a gift from James Bradner & Behnam Nabet (Addgene plasmid # 91798). Plasmids created as part of this study are available from Addgene: https://www.addgene.org/Brian_Hendrich/.

To create Auxin-depletable cell lines, parent lines were first transfected with a PiggyBac construct to constitutively express OsTir1 (created using pMGS56 (GFP-ARF16-PB1-P2A-OsTIR1); a gift from Michael Guertin (Addgene plasmid # 129668)) linked to either G418 or Hygromycin resistance, and cells were cultured under selection to maintain OsTir1 expression. To create degron knock-ins, cells were lipofected with a targeting vector and appropriate gRNA (Table 1) in a Cas9-expression vector (pSpCas9(BB)-2A-GFP (PX458); a gift from Feng Zhang; Addgene plasmid # 48138). Drug resistant colonies were genotyped and correctly targeted clones were subsequently transiently transfected with an expression plasmid for Dre recombinase to remove ROXed drug selection cassettes. *Chd4* targeting required two rounds of transfection, selection, and drug removal, while *Mbd3* and *Sall4* targeting required one round. For protein depletion, cells were treated with 500 µM Auxin or 500 µM dTAG-13 in standard culture media. All HALOTag-fusion ES cell lines were heterozygous for the HALOTag fusion, i.e. HALO/+, and verified by western blotting.

**Table 1.**
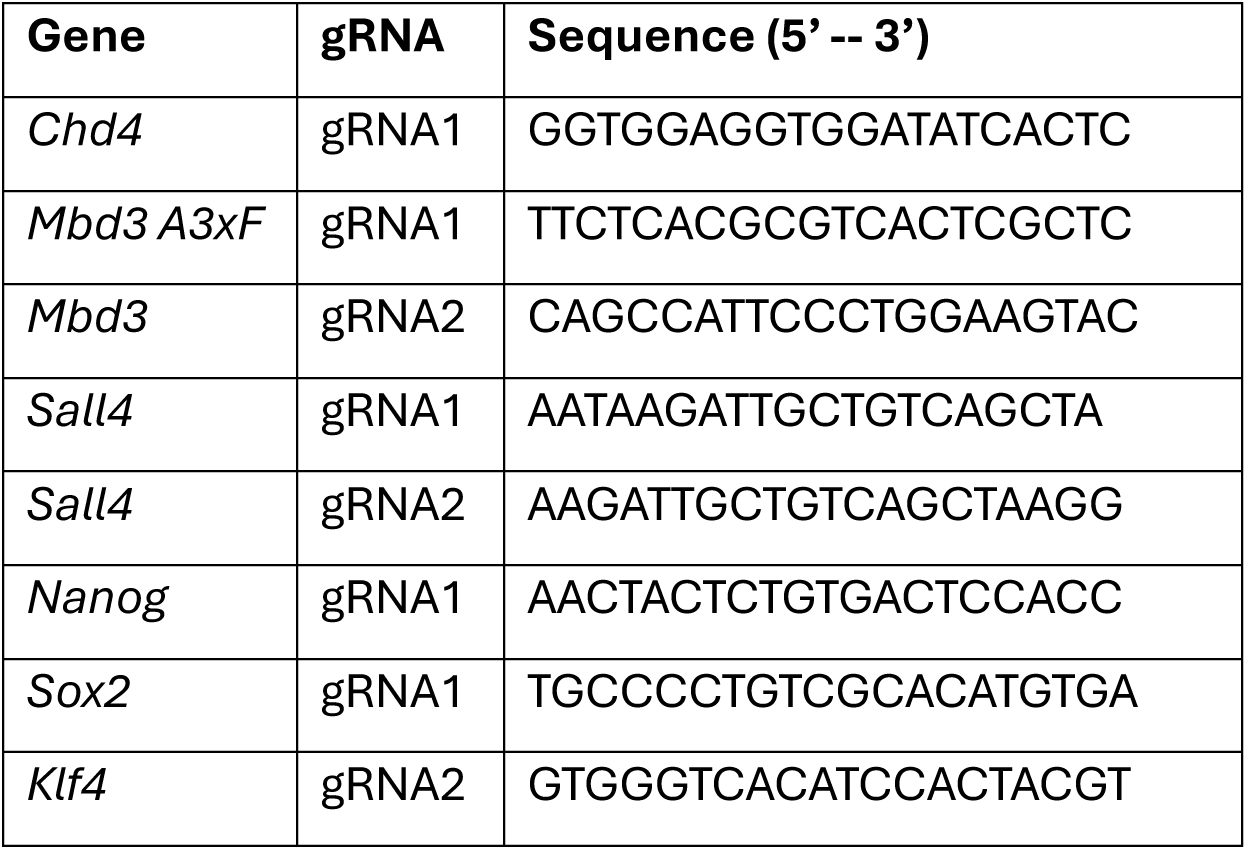
gRNA sequences used for gene targeting.

### Nuclear extract fractionation and western blotting

Nuclear fractionation was carried out as described (Gillotin, 2018). Briefly, cells were collected in ice-cold PBS and pelleted in a refrigerated centrifuge. The cell pellet was lysed by gentle up and down pipetting five times in ice-cold buffer E1 (50 mM HEPES, 140 mM NaCl, 1 mM EDTA, 10% glycerol, 0.5% NP40, 0.25% Triton X-100, 1 mM DTT, protease inhibitors). After pelleting and washing in E1 buffer, the pellet was resuspended in ice-cold E2 buffer (10 mM Tris-HCl, 200 mM NaCl, 1 mM EDTA, 0.5 mM EGTA, protease inhibitors) and shaken for 45 minutes at 1400rpm at 4°C. The supernatant, representing the nuclear fraction, was collected into a fresh tube. After washing in E2 buffer, the pellet was resuspended in ice-cold E3 buffer (50 mM Tris-HCl, 20 mM NaCl, 1 mM MgCl2, 1% NP-40, protease inhibitors). The resuspended pellet was sonicated at 4°C in a Bioruptor Plus (Diagenode) for 5 minutes, using 30-second ON/30-second OFF cycles at high power. Following sonication, nuclear and chromatin fractions were centrifuged at 16,000xg at 4°C for 10 minutes. 10 µg of extract per lane was used for western blots. Antibodies are listed in Table 2.

**Table 2.** Antibodies used in this study with dilutions used for western blot.

| Antibody target | Raised in | Supplier | Cat # | Dilution (WB) |
| --- | --- | --- | --- | --- |
| CHD4 | Mouse | Abcam | ab70469 | 1/5000 |
| CHD4 | Rabbit | Abcam | ab72418 | 1/10000 |
| FLAG | Mouse | Sigma Aldrich | F3165/F1804 | 1/2000 |
| HA | Mouse | Invitrogen | 26183 | 1/2000 |
| ADNP | Goat | R&D<br>Systems | AF5919 | 1/2000 |
| Histone H3 | Rabbit | Abcam | ab1791 | 1/5000 |
| MTA2 | Mouse | Abcam | ab50209 | 1/5000 |
| NANOG | Rabbit | Bethyl<br>Labs | A300-387A | 1/2000 |
| SOX2 | Rat | E-bioscience | 14-9811-82 | 1/2000 |
| Lamin B1 | Rabbit | Abcam | ab133741 | 1/10000 |
| Mouse IgG | Rabbit | Invitrogen | A16162 | 1/100 |
| Rabbit IgG | Guinea<br>Pig | Antibodies<br>Online | ABIN101961 | 1/100 |
| Goat IgG | Rabbit | Abcam | ab6697 | 1/100 |
| Rat IgG | Rabbit | Abcam | ab6703 | 1/100 |
| Mouse IgG | Sheep | VWR | NA931V | 1/2000 |
| Rabbit IgG | Donkey | VWR | NA934V | 1/2000 |

### Cell cycle analysis

The cell cycle distributions of auxin (500 μM) or dTAG-13 (500 nM) treated ES cell lines were assessed using propidium iodide (PI) staining coupled with flow cytometry. Approximately 200,000 single cells were fixed using 70% ethanol for at least 24 hours before staining. Fixed samples were washed twice in PBS and resuspended in 200 μL of PI solution [50 μg/mL propidium iodide (Invitrogen, P3566) and 50 μg/mL DNase and protease-free RNase A (ThermoScientific, EN0531), diluted in sterile PBS and incubated overnight at 4°C in the dark. The fluorescent intensity of stained samples was determined using the Attune NxT (Thermofisher) equipped with a 561 nm laser line and a minimum of 10,000 single cell events were recorded. The resulting FCS files were analysed using FlowJo (v10.10) and tested for significance using a mixed-effects model with Dunnett’s multiple comparisons correction.

### Single molecule imaging

ES cells were passaged 24 hours before imaging onto either: No 1.0 35 mm glass bottom dishes (MatTek Corporation P35G-1.0-14-C) or No 1.5 35 mm glass bottom dishes (MatTek Corporation P35G-1.5-14-C) with their surfaces pre-coated in poly-L-ornithine (Sigma Aldrich P4957) for ≥30 minutes at 37°C, followed by three PBS rinses at room temperature, followed by 100<u>µg/ml</u> Laminin (Sigma Aldrich L2020) coating in PBS for >4 hours at room temperature. Cells were labelled on the day of imaging with either 250 nM HaloTag®-PA-JF646 (a gift from L. Lavis, Janelia) for 15 minutes, rinsed twice in PBS and incubated for 20 minutes at 37°C in fresh media. PBS rinsing and 20-minute incubation steps were repeated five more times, and the cells were then imaged in fresh media.

Single-molecule tracking was carried out using oblique illumination (Tokunaga et al., 2008) on a custom-built double helix point spread function microscope with a Nikon Eclipse Ti-U inverted microscope body and a box incubator set to 37°C (Carr et al., 2017). Beams were expanded and collimated using Galilean beam expanders, then combined using dichroic mirrors. For 2D imaging, a Nikon 1.49 NA 60× oil immersion objective (CFI Apochromat TIRF 60XC Oil) was used to focus excitation beams onto the sample, and the microscope was also set to an internal magnification of 1.5x. For 3D DHPSF imaging, a Nikon 1.27 NA 60x water immersion objective lens (CFI SR Plan Apo IR 60XC WI) was used without internal 1.5x magnification. The emission path of the microscope was also modified to include a fixed double helix phase mask (DoubleHelix, Boulder, CO) in the Fourier domain of the emission path of the microscope (Carr et al., 2017; Pavani et al., 2009). For long-exposure imaging, control samples were fixed in 4% formaldehyde (v/w) in PBS for 10 minutes at room temperature, rinsed twice with PBS and stored in PBS. Samples were imaged in identical pre-warmed culture media and under identical conditions to live-cell samples. Across all experiments, three separate samples per condition were imaged in a day and all experiments were repeated on at least two separate days for biological replications.

Background subtraction with a five-pixel rolling ball radius was carried out on image stacks using open-source software Fiji (Schindelin et al., 2012). Single molecules imaged in 2D were localised using the PeakFit tool within the GDSC Single Molecule Light Microscopy (SMLM) plugin (https://github.com/aherbert/gdsc-smlm (Etheridge et al., 2022)) for Fiji. Localisations were filtered for precision better than 25 nm. For single molecules imaged in 3D, PeakFit was used as above but with an initial precision threshold of 40 nm and an additional analysis step after this. 2D localisations of DHPSF lobes were paired to generate 3D localisations using DHPSFU (https://github.com/TheLaueLab/DHPSFU). Localisations were then tracked across frames to generate trajectories using custom Python code (https://github.com/wb104/trajectory-analysis). Localisations were connected between two successive frames if they were located within 400 nm of each other.

Short-exposure trajectories were segmented into confined and unconfined sub-trajectories using the 4P-algorithm from (Basu et al., 2023). Bound fractions were calculated for a sample based on the number of trajectory frames assigned as being confined, divided by the total number of trajectory frames. Tracks with a residence time shorter than 1.5 seconds were filtered out to reduce the impact of noise on further analysis. The decay curve of residence times for each sample was fitted to a single exponential fit using MATLAB (2018) to yield apparent dissociation rates. The measured apparent dissociation rate (*k*_*off*_^*app*^) was calculated by fitting a single exponential decay function to the survival (Kaplan-Meier) function (*S*) of TF dwell times measured from the point of first observation (*t*), as below:

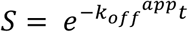

### RNAseq

Cells were harvested in Trizol Reagent and RNA purified using Zymo Direct-zol columns (Zymo Research) according to manufacturer’s instructions. 60-100 ng of Ribosomal RNA-depleted mRNA was used for library preparation with the the NEXTflex RapidDirectional RNA-seq kit (Illumina). 100 bp paired end (2 x 50 bp) sequencing was performed on a NovaSeq S1 flowcell at the CRUK Cambridge Institute sequencing facility.

Raw data were trimmed using TrimGalore v0.6.4. Reads were indexed and aligned to the unmasked GRC38/mm10 reference genome using the Burrows-Wheeler Aligner (BWA) (Li and Durbin, 2009). Successful removal of adapter and low-quality bases was assessed using FastQC after trimming. File conversion, sorting, removal of duplicates and mitochondrial reads were performed using samtools (Li et al., 2009).

Read counts for genomic features were summarised using featureCounts (Liao et al., 2014) and a list of the GRC38/mm10 genomic features in a GTF format downloaded from https://www.ensembl.org/ and https://genome.ucsc.edu/cgi-bin/hgTables. Differential expression was analysed using DESeq2 (Love et al., 2014). Raw expression counts were transformed and normalised to FPKM (fragments per kilobase of transcript per million mapped reads). Simple linear models were used for pairwise comparative analyses between the 0-timepoint and depletion timepoints. An adjusted p-value threshold of 0.05 was used to identify significantly differentially expressed genes. For principal component analysis (PCA) and the generation of correlation matrices vst-transformed data were used within DESeq2.

Genes were labelled with their Ensembl and their common gene symbols using the Annotation Hub and ensembldb packages (Rainer et al., 2019). The ComplexHeatmap package was used for the generation of expression heatmaps (Gu et al., 2016). Volcano plots were generated using the EnhancedVolcano package. To find enriched gene ontology terms among differentially expressed genes, the enrichGO function of the clusterProfiler R package and the genome-wide annotations were retrieved using the biomaRt package (Yu et al., 2012).

### ATAC-seq

100,000 nuclei were used for each reaction using the OMNI-ATAC protocol (Corces et al., 2017) with 10,000 rat nuclei added as a spike-in control. Tagmentation was achieved using Tn5 made in-house (Picelli et al., 2014) for 30 minutes at 37°C on a shaking block. Immediately after incubation DNA was purified using a Zymo DNA Clean and Concentrator Kit (Zymo Research). DNA was then eluted in 21 μL of DNAse/RNAse-free H2O. Barcoding for library preparation was performed by PCR amplification using NEBNext® High-Fidelity 2X PCR Master Mix (New England Biolabs) and NEBNext® index primers for Illumina sequencing. Amplified samples were purified using AMPure SPRI beads (VWR International Ltd) and resuspended in 25 μL of 10mM Tris-HCl pH 8.0. The Agilent 4200 TapeStation System with D1000 ScreenTape and D1000 Reagents (Agilent) was used for quantification and fragmentation check of the samples. Equimolar ratios of all samples (7 conditions x 2 biological replicates x 2 technical replicates each) were pooled for 300 bp paired end (2 x150 bp) sequencing performed on a NovaSeq S2 flowcell with the CRUK Cancer Institute sequencing facility.

Raw data were trimmed to remove adapter contamination using TrimGalore v0.6.4. Reads were aligned to the GRC38/mm10 reference genome and the Rattus norvegicus/Rnor_6.0 genome (spike-in) using the BWA aligner. File conversion, sorting, removal of duplicates and mitochondrial reads were performed using SAMtools (Li et al., 2009). Fragments up to 120 bp in length, which represent nucleosome free regions, were used for further analysis. For visualisation purposes bigwig files were generated using bamCoverage from deepTools (Ramírez et al., 2014). A scale factor was applied calculated as the number of uniquely mapped rat spike-in reads for each sample divided by the number of uniquely mapped reads in the sample with the lowest count. All replicates were pooled together into a single track using bigWigMerge from UCSC Tools (Kent et al., 2002). Mean profiles of the signal and heatmaps of these regions were plotted using the ComputeMatrix, plotHeatmap and plotProfile functions from deepTools.

Peak calling was performed for ≤120bp fragments using macs2 (Zhang et al., 2008) and applying the -f BAMPE -q 0.05 –nolambda –keep-dup auto parameters. Peak annotation and motif analysis was conducted using the annotatePeaks and findMotifsGenome functions from HOMER (Heinz et al., 2010). Integration of these annotated peaks with RNA-seq gene expression data was conducted using custom R scripts. Differential accessibility analysis was carried out using DiffBind (https://bioconductor.org/packages/DiffBind). Differentially accessible regions were identified in pairwise comparisons to timepoint-0 using the rat aligned reads as a spike-in and an adjusted p-value threshold of 0.05. Chromatin state enrichment analysis for both ADNP and CHD4 differentially accessible regions was performed using ChromHMM (Ernst and Kellis, 2017), using a predefined chromatin state model generated from E14 mouse ESCs ChIP-seq data (Pintacuda et al., 2017).

To analyse the Tn5 integration sites of the ATAC-seq data the plotFootprint() function in VplotR (Serizay and Ahringer, 2021) was used with the addition of a code in a loop to calculate the normalised version of the cuts by manually dividing by the library size of each merged bam file. To analyse the fragment sizes of the ATAC-seq data that correspond to different nucleosome structure sizes, the plotVmat() function in VplotR was used. The merged reads of different time points of CHD4 depletion were normalised using the native libdepth+nloci option, which normalises the reads using the library depth and the number of loci.

### CutGRun and CutGTag

CutCRun and CutCTag were carried out as described (Janssens et al., 2022; Meers et al., 2019). For CutCRun 100,000 live ES cells were used per reaction, with 10,000 rat nuclei added as a spike-in control. CutCTag was performed on 100,000 nuclei per reaction. Antibodies (Table 2) were used at 1/100 dilution. pAG-MNAse and pA-Tn5 were made and purified in-house as described (Kaya-Okur et al., 2019; Meers et al., 2019). Plasmids 3XFlag-pA-Tn5-Fl and pAG/MNase were a gift from Steven Henikoff (Addgene plasmids #124601 and #123461, respectively). Library preparation for sequencing was carried out in the CSCI Genomics facility. 300 bp paired-end sequencing was performed on a NovaSeqX 25B flowcell at the CRUK Cancer Institute sequencing facility.

Raw data were trimmed to remove adapter contamination using TrimGalore v0.6.4. Reads were aligned to the unmasked GRC38/mm10 reference genome and the Rattus norvegicus/Rnor_6.0 genome (spike-in) using the BWA. File conversion, sorting, removal of duplicates and mitochondrial reads were performed using SAMtools. For visualisation purposes bigwig files were generated using bamCoverage from deepTools. A scale factor was applied calculated as the number of uniquely mapped rat spike-in reads for each sample divided by the number of uniquely mapped reads in the sample with the lowest count. All replicates were pooled together into a single track using bigWigMerge from UCSC Tools. Mean profiles of the signal and heatmaps of these regions were plotted from these tracks using the ComputeMatrix, plotHeatmap and plotProfile functions from deepTools. Peak calling was performed using macs2 and SEACR (Meers et al., 2019). SEACR was used in non-control mode with the stringent threshold setting (threshold = 0.01). Input BED files consisted of scaled fragment bedgraphs generated from CUTCRUN data. Scaling was performed using a factor based on the number of uniquely mapped rat spike-in reads for each sample, which was then adjusted relative to the sample with the fewest mapped reads. Motif analysis was conducted using HOMER. Differential accessibility analysis was also carried out using the DiffBind package, incorporating DESeq2 functionality. Differentially bound peaks were identified in pairwise comparisons to timepoint-0 using an adjusted p-value threshold of 0.05. Chromatin state enrichment analysis for differentially bound regions was performed using ChromHMM as for ATAC-seq.

### MNase-seq

All MNase experiments, sequencing and data processing were carried out exactly as described (Bornelöv et al., 2018) except that nuclei were digested with 500 U/ml micrococcal nuclease (New England Biolabs) at 24°C for 15 minutes with shaking.

## Accession numbers

ATAC-seq: E-MTAB-15037

RNAseq: E-MTAB-15102

nascent RNAseq: E-MTAB-15127

CutCRun: E-MTAB-15606, E-MTAB-15607

CutCTag: E-MTAB-15625

Sall4 Depletion ATAC-seq: E-MTAB-15375

Sall4 CutCRun: GSE203303 (Ru et al., 2022)

MNase: E-MTAB-6807 (Bornelöv et al., 2018), PRJNA1332303

## Acknowledgments

We are grateful to Nick Owens, Vladimir Teif, Joel Mackay and past and present Hendrich and Laue group members for helpful discussions; to David Klenerman and Ziwei Zhang for help with imaging; to Marco Trizzino for support and to the CSCI facilities for expert technical assistance.

## Author Contributions

AK, OO, IMB, DS and BH created and validated ES cell lines; AK, OO and NR generated sequencing data; AK, OO and RR analysed sequencing data; DS and EDL conducted single molecule imaging experiments and analyses; ML devised methodology; NR, RR, EDL and BH supervised the project and EDL and BH obtained funding. The manuscript was written by BH with input from all other authors.

---

**Supplemental Figure 1.**
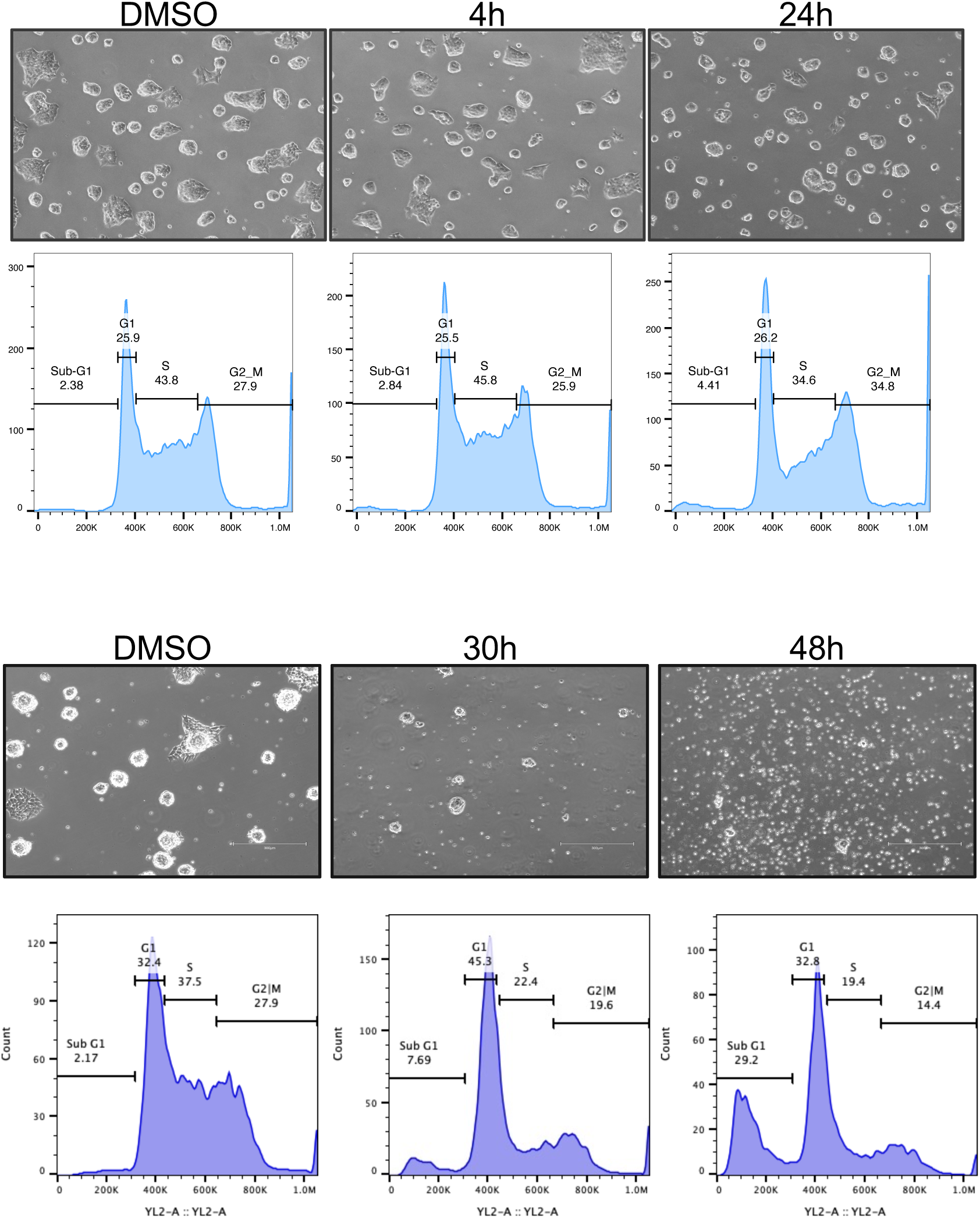
Cell cycle arrest and cell death following CHD4 depletion. Phase contrast images and example flow cytometry plots of CHD4-mAID ES cells in 2iL conditions after indicated times of Auxin addition. Cell cycle stage and relative percentage of cells are indicated on the flow cytometry plots.

**Supplemental Figure 2.**
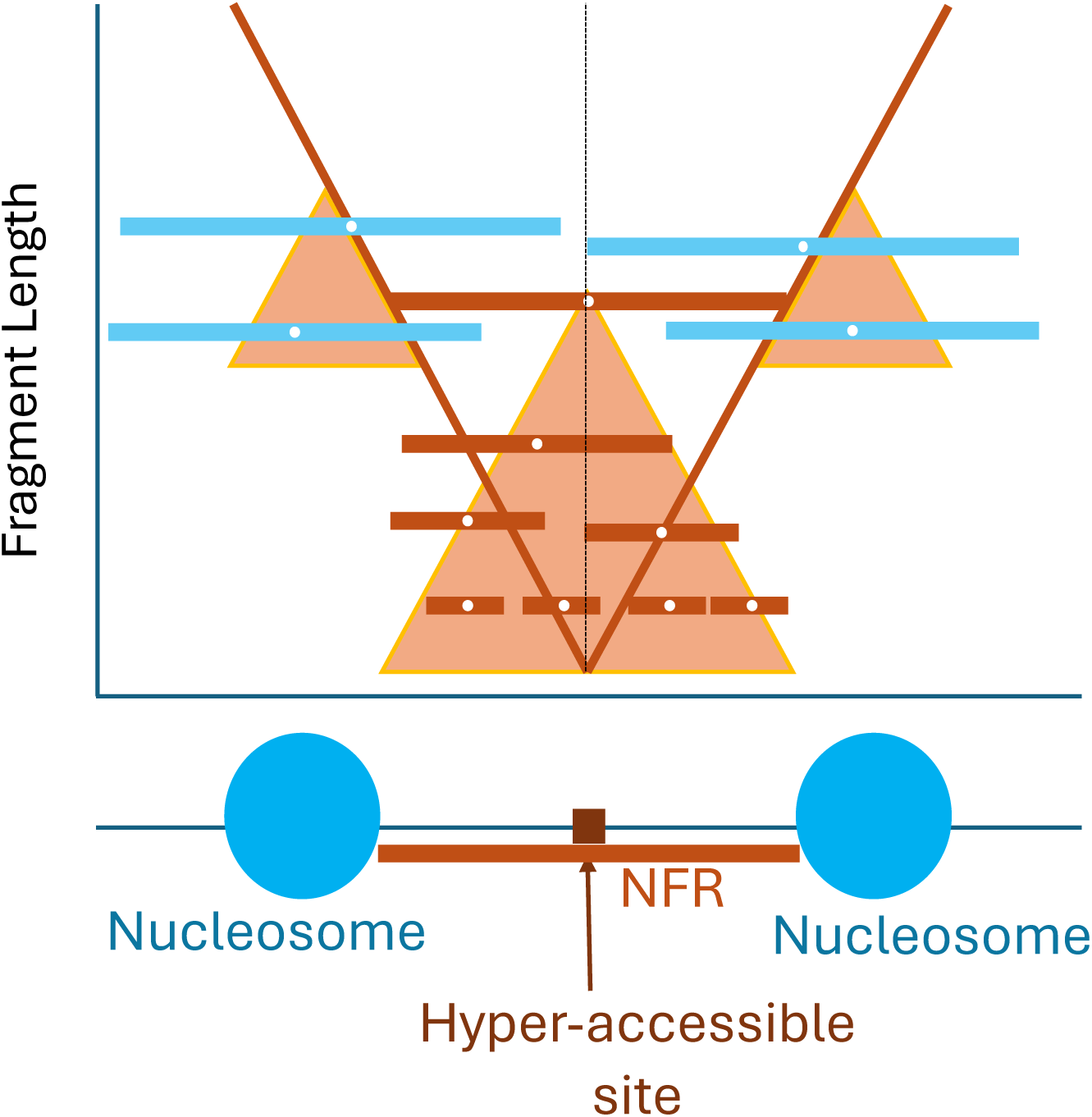
Schematic diagram of Vplots. Adapted from (Henikoff et al., 2011). The Vplot is derived by plotting the midpoint of all recovered fragments (horizontal lines) onto the graph, with midpoint position (white circle) on the x-axis and fragment length on the y-axis. Red lines correspond to reads obtained from fragments with both ends (i.e. Tn5 integration sites) in the NFR, while blue lines represent reads spanning a nucleosome. The vertical dotted line indicates the position of the central hyperaccessible site. The inferred chromatin structure of the locus is shown below.

